# Bacterial Infection of the Placenta Induces Sex-Specific Responses in the Fetal Brain

**DOI:** 10.1101/2022.05.03.490463

**Authors:** Kun Ho Lee, Matti Kiupel, Thomas Woods, Prachee Pingle, Jonathan Hardy

## Abstract

**BACKGROUND:** Epidemiological data indicate that prenatal infection is associated with an increased risk of several neurodevelopmental disorders in the progeny. These disorders display sex differences in presentation. The role of the placenta, which is a target of prenatal infection, in the sex-specificity of neurodevelopmental abnormalities is unknown. We used an imaging-based animal model of the bacterial pathogen *Listeria monocytogenes* to identify sex-specific effects of placental infection on neurodevelopment of the fetus.

**METHODS:** Pregnant CD1 mice were infected with a bioluminescent strain of *Listeria* on embryonic day 14.5 (E14.5). Excised fetuses were imaged on E18.5 to identify the infected placentas. The associated fetal brains were analyzed for gene expression and altered brain structure due to infection. The behavior of adult offspring affected by prenatal *Listeria* infection was analyzed.

**RESULTS:** Placental infection induced sex-specific alteration of gene expression patterns in the fetal brain and resulted in abnormal cortical development correlated with placental infection levels. Furthermore, male offspring exhibited abnormal social interaction, whereas females exhibited elevated anxiety.

**CONCLUSION:** Placental infection by *Listeria* induced sex-specific abnormalities in neurodevelopment of the fetus. Prenatal infection also affected the behavior of the offspring in a sex-specific manner.

**Impact:**

- Placental infection with *Listeria monocytogenes* induces sexually dichotomous gene expression patterns in the fetal brain.
- Abnormal cortical lamination is correlated with placental infection levels.
- Placental infection results in autism related behavior in male offspring and heightened anxiety level in female offspring.

## INTRODUCTION

The molecular and cellular mechanisms leading to most neuropsychiatric disorders, such as autism spectrum disorder (ASD), remain unclear due to their complex polygenic etiology. Epidemiological data indicate that prenatal infection with bacterial, viral, or parasitic pathogens during pregnancy is associated with an increased risk of neuropsychiatric disorders in the progeny, including ASD^1^ and schizophrenia^2, 3^. Injection of bacterial endotoxin lipopolysaccharide (LPS) or polyinosinic-polycytidylic acid [poly(I:C)], which mimics viral infections, activates the immune system of pregnant rodents and results in altered brain gene expression^4^ and atypical behavior in offspring^5^. These behavioral abnormalities are notably relevant to ASD core symptoms, such as repetitive behaviors and deficit in social interaction. Furthermore, animal studies show sex biased behaviors and responses in offspring after exposing to LPS and poly(I:C) during pregnancy, which resembles sex differences in neuropsychiatric disorders, including ASD^6, 7^. Maternal immune activation (MIA) induced by LPS or poly(I:C) causes the changes in fetal brain development. Although injection of immunogens in pregnant animals results in consistent altered behavior and brain abnormalities in the progeny, they do not elicit the complex immune responses induced by actual infection. Prenatal pathogens exhibit tissue and cell-specificity as well as directed immune modulation, such that the different pathogens may regulate MIA differently. For example, infection of rats with Group B *Streptococcus* elicits distinct MIA patterns including neutrophil infiltrates that differ from immune stimulants such as LPS and poly(I:C)^8^. In addition, prenatal influenza is a risk factor for schizophrenia^2^, whereas no such association was found with prenatal infection with either maternal type 1 herpes simplex virus^9, 10^ or cytomegalovirus^11^. Thus, the induction of MIA is complex and cannot be completely replicated by any single approach. It is therefore critically important to examine different prenatal infection models and their specific effects on fetal brain development and behavior.

*Listeria monocytogenes* (*Lm*) provides an excellent animal model for prenatal infection^12, 13^. This Gram-positive bacterium is a foodborne pathogen and is a significant health concern during pregnancy because pregnant women are up to 10 times more likely to be infected with *Lm*^14^. An important hallmark of prenatal listeriosis is the infection of the placenta^15–17^. Placental infection by *Lm* can lead to many overt fetal and newborn pathologies, including spontaneous abortions, stillbirth, and other neonatal illnesses, even while pregnant mothers can be largely asymptomatic^12, 18–20^. Previously, we reported that bradycardia was only observed in fetuses with infected placentas within the same infected pregnant mouse as those with normal heart rates^21^. These studies demonstrated that bradycardia induced by placental infection was not systemic but localized.

Infection with the appropriate dose of intravenous *Lm* on embryonic day 14.5 results in abortion, stillbirth, and fetal bradycardia in the absence of overt maternal disease symptoms. Although placental infection by *Lm* causes adverse outcomes in newborns, neurodevelopmental consequences of this infection have not been characterized. In addition, sex-specific consequences of placental infection have not been defined for any living pathogen. The aims of this study were to understand how bacterial infection of the placenta affects fetal neurodevelopment, to determine if sex-specific responses occur, and to assess effects on the behavior of the offspring.

## METHODS

### Animal care and use

All animal procedures were approved by the Institutional Animal Care and Use Committee and the Biosafety of Michigan State University under protocol number 201800030. Michigan State University (MSU) has approved Animal Welfare Assurance (A3955-01) from the NIH Office of Laboratory Animal Welfare (OLAW). In addition, all components of the University are accredited by the Association for Assessment and Accreditation of Laboratory Animal Care, International (AAALAC Unit #1047). Standard BSL-2 containment and handling procedures were used for all animals including the offspring. These procedures were also approved according to the specific MSU Biosafety Protocol 0000058, and all laboratories, procedure rooms and facilities are inspected by MSU Environmental Health and Safety. Timed CD1 pregnant mice purchased from Charles River Laboratories were used for all studies and housed in temperature controlled, 12:12 hour light and dark cycle rooms. Euthanasia was performed by cervical dislocation under isoflurane anesthesia by trained personnel according to NIH and MSU approved protocols.

### *In vivo* bioluminescence imaging (BLI) and tissue processing

The bioluminescent strain of *L. monocytogenes* used in this study (Perkin Elmer Xen32) was generated in a 10403S strain background^22^. Cultures were incubated overnight at 37°C in brain heart infusion (BHI) broth. The overnight culture was sub-cultured in fresh BHI broth to an optical density (OD_600_) of 0.5. Timed embryonic day 11 (E11) pregnant CD1 mice were house at the Michigan State University Clinical Center animal facility under BSL-2 containment. Pregnant mice were administered a tail vein injection of 2 x 10^5^ colony-forming units (CFU) of Xen32, diluted in 200 uL phosphate-buffered saline (PBS), or an equivalent volume of PBS vehicle on E14.5. On E18.5, pregnant mice were anesthetized using isoflurane and imaged using the *In vivo* bioluminescence imagining system (IVIS; Perkin Elmer Inc.), and then humanely scarified by cervical dislocation while under isoflurane anesthesia according to the approved animal protocol. Uterine horns were excised immediately and imaged again using the IVIS. Individual fetuses could be imaged separately for high-resolution BLI. Signal levels from the placenta at this dose and timing vary over orders of magnitude within one pregnant mouse, permitting the analysis of systemic versus localized effects and allowing for comparisons of fetal brains with and without high placental BLI signal from the same pregnant animal. Fetal brains were collected and transferred into sterile Eppendorf tubes, flash-frozen, and stored at -80°C until analyzed.

### Histology

For histology of the fetal brain, excised fetuses and placentas were imaged with ex vivo BLI to determine signal levels of the associated placentas. The heads were removed and fixed overnight in 4% paraformaldehyde for sectioning. Following sectioning, the brains were routinely processed and embedded in paraffin and matched coronal sections were stained with hematoxylin and eosin (H&E). Matched coronal sections were also obtained from PBS-injected pregnant mice and from fetuses with and without detectable BLI signals from the placenta from infected pregnant mice. BLI signals from the placenta were measured with identical regions of interest (ROIs). For immunohistochemistry of brains of the adult offspring, the animals were euthanized with CO_2_ according to the approved animal protocol. The brains were removed, fixed in 4% paraformaldehyde and matched coronal sections were obtained as described above. Several sections from each brain were stained with H&E following the same routine methods or immunohistochemically labeled with a rabbit monoclonal anti-c-Fos antibody (dilution 1:1,000, EPR21930-238, Abcam, Boston, MA). Immunohistochemistry was performed on the Dako link 48 Automated Staining System (Agilent Technologies, Santa Clara, CA) using a high pH antigen retrieval and peroxidase-conjugated EnVision Polymer Detection System (Aligent Technologies) with 3,3’-diaminobenzidine (DAB) as the chromogen and hematoxylin counterstaining.

### RNA-seq

Total RNA was isolated using the phenol/guanidine based QIAzol lysis reagent (Qiagen, Valencia, CA), according to the manufacturer’s recommendations. The concentration and quality of RNA samples were measured using Qubit (ThermoFisher) and BioAnalyzer (Agilent), respectively. Samples with RNA integrity number values of 9 or above were selected for sequencing. Fetal brains (positive BLI signal n = 19; control n =6) were collected and total RNA was submitted for next generation sequencing (NGS) library preparation and sequencing to Research Technology Support Facility at Michigan State University. Libraries were prepared using the Illumina TruSeq Standard mRNA Library Preparation Kit with IDT for Illumina Unique Dual Index adapters following manufacturer’s recommendations. Completed libraries were quality checked and quantified using a combination of Qubit dsDNA High Sensitivity and Agilent 420 TapeStation HS DNA1000 assays. Libraries were pooled in equimolar proportions for multiplexed sequencing. The pool was quantified using the Kapa Biosystems Illumina Library Quantification qPCR kit. This pool was loaded onto two lanes of an Illumina HiSeq 4000 flow cell (two technical replicates) and sequencing was performed in a 1 x 50 single read format using HiSeq 4000 SBS reagents. Base calling was done by Illumina Real Time Analysis v2.7.7 and output of RTA was demultiplexed and converted to FastQ format with Illumina Bcl2fastq v2.19.1. The raw single-end (SE) reads were processed to trim sequencing adapter and low-quality bases. The clean SE RAN-seq reads were mapped to the mouse reference genome (GCRm38.p6/mm10) using STAR (Spliced Transcriptions Alignment to a Reference) v2.3.2^23^. Mapped reads were assigned to genes with FeatureCounts in the subread package^24^.

### Differential gene expression analysis and functional enrichment analysis

Differential gene expression analysis was performed using DESeq2 v1.32.0^25^ in R v4.1.1. Genes with minimum 5 raw reads in at least 20 samples were filtered out, resulting a total of 19,180 of genes. Differentially expressed genes with p-adj < 0.05 were used to perform functional enrichment analysis using the g:Profiler system (https://biit.cs.ut.ee/gprofiler/gost)^26^. Biological pathways with p-adj < 0.05 were considered significant. To examine the sex dependent effects of placental infection, a female specific gene, *Xist*, was used to identify the sex of fetal brains from RNA-seq samples. Differential gene expression and functional enrichment analyses were performed using the same parameters as placental infection. Volcano plots were generated using EnhancedVolcano package in R^27^.

### Social interactions and repetitive behaviors

The three-chamber social approach assay has been widely used to test for assaying sociability in mice^28^. This assay measures interaction between animals that are provided choices between unfamiliar animals and inanimate objects (social interaction). We used a custom three-chamber apparatus (63 cm x 30 cm x 31 cm) with an empty middle chamber and accessible side chambers on either end that contain cylindrical open barred cages in which mice or objects are placed. An inanimate object is placed in one of the barred cages in one side chamber, and an unfamiliar mouse is placed in the barred cage in the other side chamber. A test subject mouse is placed in the central chamber and allowed to freely interact with the mice or objects in the side chambers. The social interaction test we employed had three phases. First, the test subject (prenatal *Lm*-exposed male = 10 and female = 8; control male = 5 and female = 5; 8 – 12 weeks of age) was habituated in the center of chamber for 10 minutes and two doorways in the chambers were closed. Second, the test subject was habituated to all three chambers for 10 minutes. Third, the subject was confined to the middle chamber, a novel object (lab tape) was placed in the barred cage in one side chamber, and a novel mouse (a treatment, sex, and age matched unfamiliar mouse) was placed in the other side chamber.

The social interaction in each test was recorded for 10 minutes. Sniffing time for each subject was recorded. Self-grooming, which is defined as time spent rubbing the face, scratching with a foot, or licking paws, was examined to measure repetitive and persistent behavior^28^. During the three-chamber social approach assay, self-grooming was measured by using a stopwatch.

### Open field exploration

Open field exploration tests measure anxiety, exploration and locomotion^29^. Mice (prenatal *Lm*-exposed male = 8 and female = 10; control male = 4 and female = 3, 8 – 12 weeks of age) were acclimated for 30 minutes before the assay. Mice were place in the middle of the testing area (63 cm x 60 cm x 31 cm) and underwent a 10-minute exploration period. Sessions were video recorded and analyzed using the ANY-maze Video Tracking System software.

## RESULTS

### BLI and postnatal effects of placental infection

All infections were performed by intravenous (IV) injection into pregnant CD1 mice with 5 x 10^5^ colony forming units (CFU) of the bioluminescent *Lm* strain 2C (Perkin Elmer Xen36) on E14.5. The dose and timing were selected based on our prior studies^21^. The IV route of *Lm-*infection in pregnant mice is employed rather than oral infection for several reasons; the foremost being that oral infection requires over 10^11^ CFU in CD1 mice and is not reproducible between laboratories. In contrast, IV infection is highly reproducible and bypasses the intestine, seeding the placenta directly in a dose-dependent manner. The selected dose results in stillbirth, abortion, and developmental abnormality, resembling listeriosis in pregnant women. An IVIS image of *Lm-*infected pregnant CD1 mouse is shown in Figure 1a, with BLI signals indicating different infection sites, including gallbladder, placentas, and fetuses. BLI of excised uterine horns shows that there is a range of infection severity indicated by the intensity of the BLI signals from the placentas (Fig. 1b). In addition, BLI demonstrated that *Lm-*infection was much greater in the placenta than the fetus (Fig. 1c), as most often at this dose the signal was only detectable in the placenta. This result was consistent with other studies that showed that fetal infection only occurs at high doses^12, 30^.

**Fig. 1.**
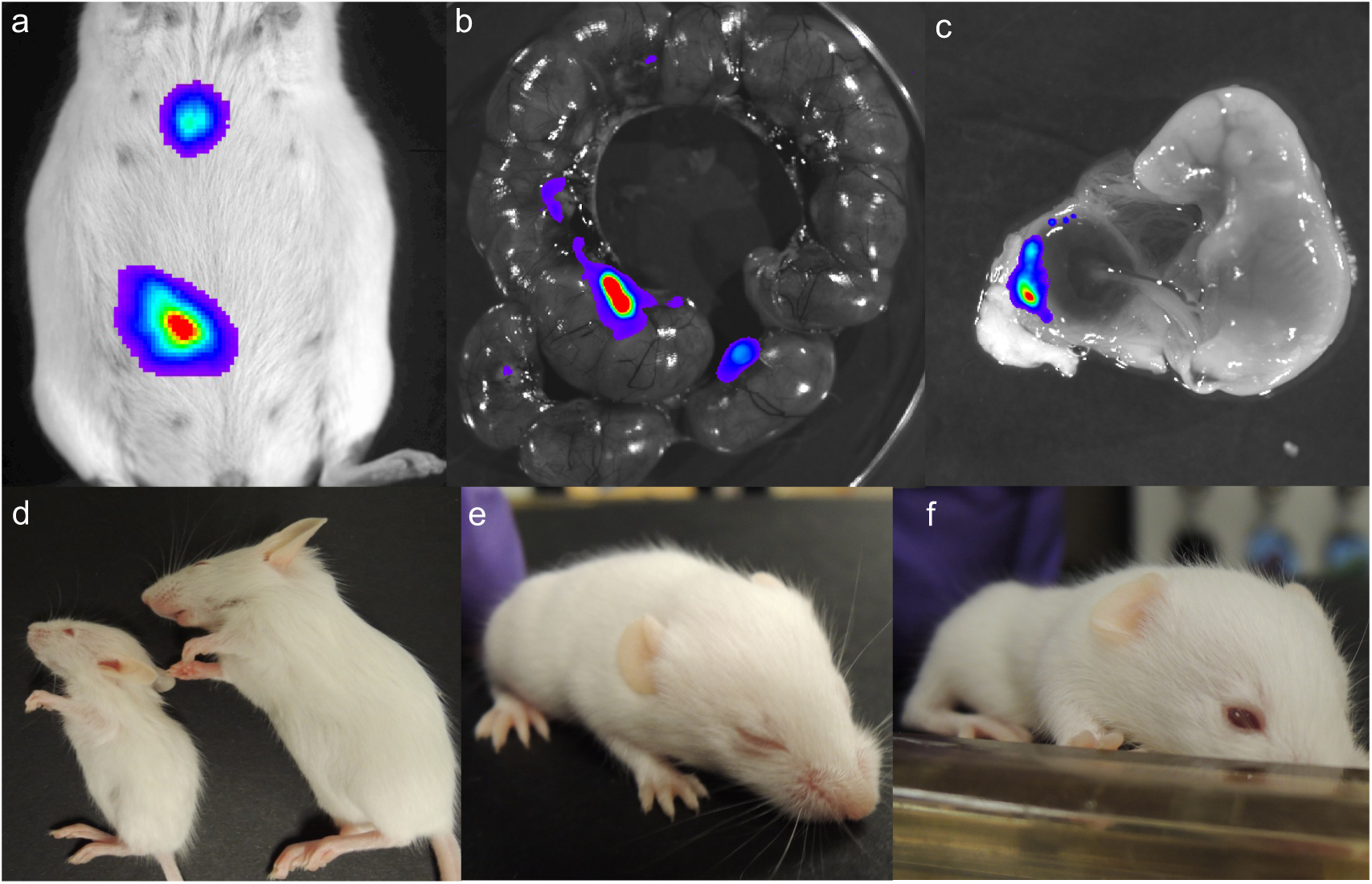
Postnatal effects of placental infection. (**a-c**) *In vivo* bioluminescence imaging (BLI) of prenatal *L. monocytogenes* (*Lm*) infection. **(a)** Live pregnant CD1 mouse on embryonic day 18.5 (E.18.5). (**b**) Excised uterine horns and **(c)** placenta. (**d**) Low birth weight due to *Lm* placental infection in a 4-week-old mouse (left) compared to a littermate (right). **(e)** *Lm*-exposed offspring exhibiting delayed eye opening compared to controls **(f)** on postnatal day 13.

When the *Lm*-infected pregnant CD1 mice gave birth, the pups showed a range of postnatal effects of placental infection. Some pups showed extreme low birth weight and altered body morphology (4 weeks old; Fig. 1d). These effects were correlated with signal levels in the live pregnant dam. Higher signal levels of >10^5^ photons/sec produced more severe effects such as stillbirth and extremely low birth weight, whereas signals <4×10^4^ photons/sec yielded litters of normal-sized pups. In addition, pups from infected pregnant dams that showed <4×10^4^ photons/sec and were indistinguishable from controls exhibited delayed eye opening (postnatal day 13; Fig. 1e). These findings show that placental infection with *Lm* affects fetal and postnatal development. The range of effects was correlated with overall signal intensities from the live pregnant animal. At the dose we employed, none of the pregnant dams exhibited overt symptoms and they were outwardly indistinguishable from PBS-injected controls. Although some pregnant dams showed BLI signals from the area of the liver and/or gallbladder, all of them survived to give birth if they were allowed to do so.

### Effect of placental infection on fetal cortical development

To determine whether placental infection promotes morphological changes in the fetal cortex, we performed hematoxylin/eosin (H&E) staining of the cortical sections of fetal brains. Pregnant CD1 mice were infected as described above and imaged to ascertain infection levels. We compared fetuses from infected and PBS-injected controls, but also fetuses within on pregnant dam that exhibited high and low signals from the placenta. The latter observation allowed us to distinguish effects due to systemic MIA from localized effects of the placenta. In the sections, layering was abnormal in the fetal brains from mice that originated from infected dams compared to controls (Fig 2a), and fetal brains from mice where the placentas exhibiting BLI signals above background showed layering alterations compared to fetuses from the same dam where the placenta had background BLI signals from the same dam (Fig. 2b).

**Fig 2.**
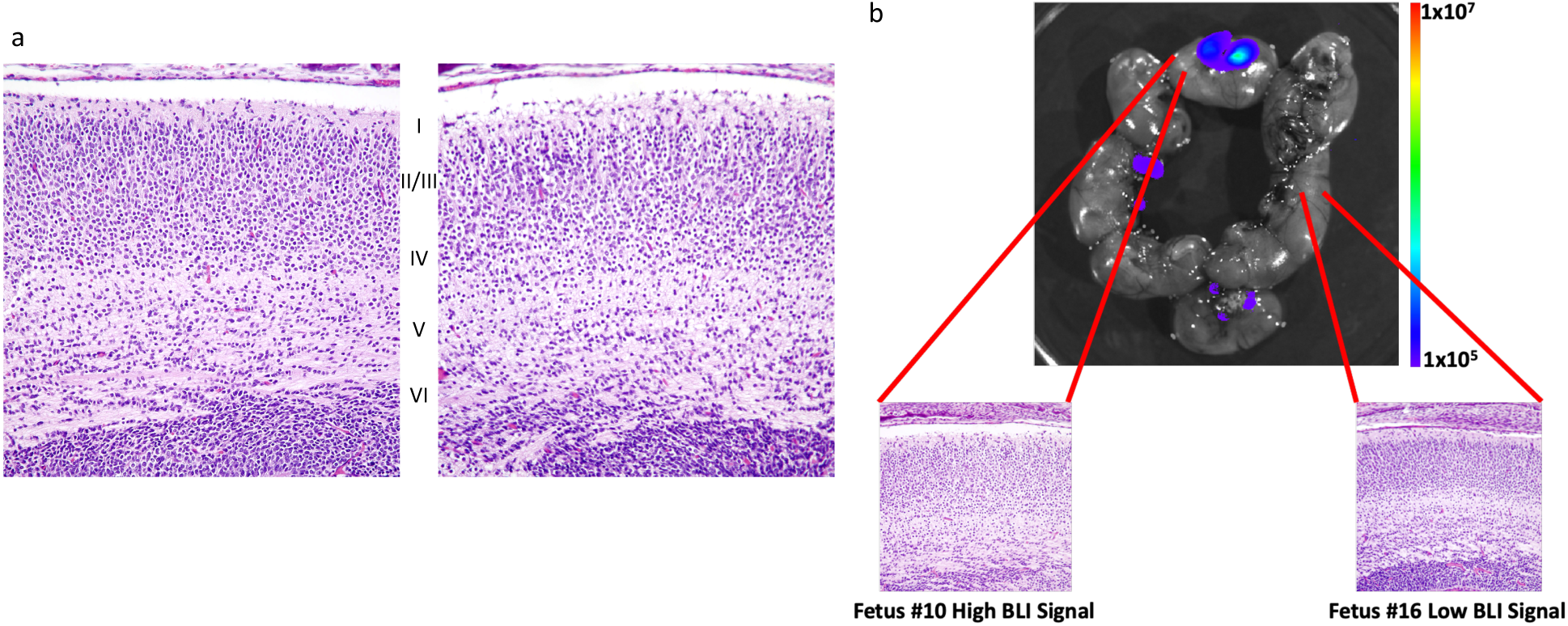
Placental infection promotes abnormal cortical lamination. **(a)** Coronal sections of fetal brain with normal layering in cortex from PBS-injected pregnant mice (left) and abnormal layering cortex from *Lm*-infected pregnant mice (right). **(b)** Abnormal layering in brains from mice with high placental BLI signal versus low placental BLI signal from the same pregnant animal. BLI signal is indicated in photons/s/cm^2^/str. Layers: I: molecular, II: external granular, III: external pyramidal, IV: internal granular, V: internal pyramidal, VI: multiform.

### Infection of the placenta alters gene expression in the fetal mouse brain

We next investigated the effect of placental infection on transcriptomic alterations in fetal brain. For these investigations, we used fetal brains from mice in which no BLI signal over background was detectable in the placenta. A total of 25 whole fetal brains (6 control and 19 *Lm-*exposed samples) were harvested on E18.5 to generate RNA-seq datasets and performed differential expression analysis using a DESeq2 package in R. The analysis revealed that IV injection of bioluminescent *Lm* into pregnant CD1 mice at E14.5 altered gene expression in the fetal mouse brain. Overall, *Lm-*exposed fetal brains had a total of 268 upregulated and 139 downregulated differentially expressed genes (DEGs) with a false discover rate (FDR) <0.05 threshold, and 1697 upregulated and 1247 downregulated with a p <0.05 threshold (Fig. 3a). Among DEGs, most significant genes include upregulated *Lyrm7*, *Flt1*, *Vegfa* and *Kdm3a*, and downregulated *Zfp125*, *Mfsd5*, *slc38a5*, *Mblac1,* and *Chd15* (Fig 3b). The Gene Ontology (GO) enrichment and KEGG analysis of upregulated DEGs revealed pathways, such as macromolecule biosynthetic and nitrogen compound metabolic processes, and hypoxia inducible factor-1 (HIF-1) signaling pathway (Fig. 3c). Furthermore, pathways such as establishment of localization in cell and protein processing in endoplasmic reticulum were identified among significantly downregulated DEGs (Fig. 3c). Many of these genes are associated with brain development or neurological function^31–34^. Together, differential expression analysis demonstrated that placental infection by *Lm* causes disruption of neurodevelopment during pregnancy.

**Figure 3.**
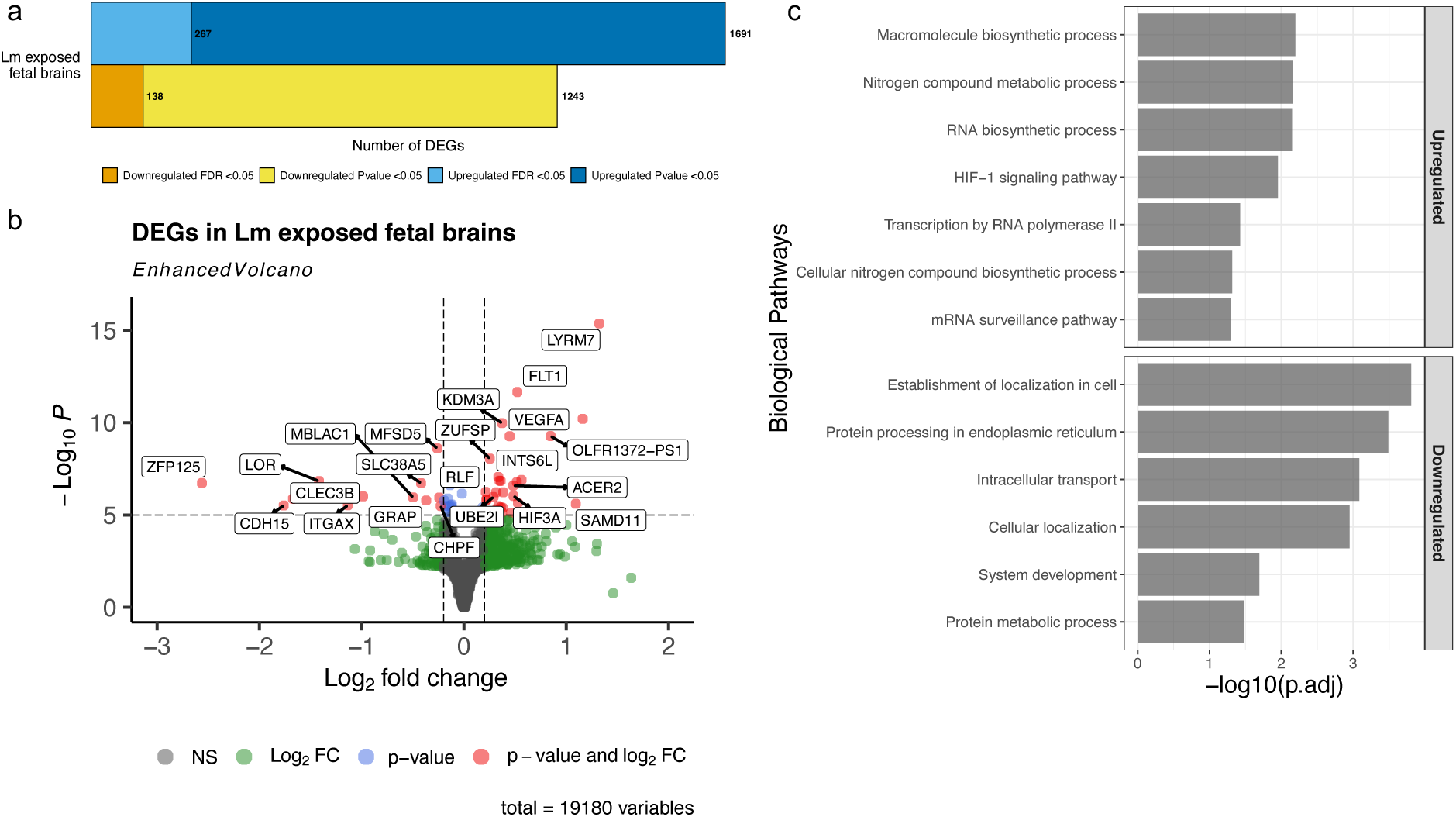
Gene expression changes in the fetal brain due to placental infection with *Lm*. (**a**) Total number of differentially expressed genes (DEGs) of fetal brains in response to placental infection. (**b**) Volcano plot of DEGs in *Lm-*exposed fetal brains of *Lm-*exposed mice at E18.5. Red dots indicate statistical significance (p-value < 10^6^) and log2(fold change) greater or less than 0.2. Total variables indicate the total number of genes that were used to generate a volcano plot. (**c**) Gene Ontology analysis of DEGs (p-adj < 0.05) in fetal brains of *Lm-*exposed mice at E18.5. Biological pathways of downregulated and upregulated fetal brains of *Lm-*exposed mice were identified using g:ProfileR.

### Male and female fetal brains exhibit distinct gene expression profiles in response to placental infection

To examine sex-specific gene expression patterns, we used *Xist*, a female specific gene, to identify sex of each fetal brain RNA-seq sample (males: 9 *Lm-*exposed and 3 controls; females: 10 *Lm-*exposed and 3 controls). We used DESeq2 in R to identify DEGs and investigated overlapping genes between both *Lm-*exposed sexes. A total of 44 and 42 downregulated DEGs were identified for males and females, respectively, with one gene overlapping between the sexes (Fig. 4a). Interestingly, females had 171 upregulated DEGs while males had 50 upregulated DEGs with 7 DEGs overlapping between the sexes (Fig. 4b). GO enrichment and KEGG analysis of upregulated DEGs of *Lm-*exposed male fetal brains identified pathways, such as VEGF receptor 2 binding, HIF-1 signaling, and microtubule organizing center (Fig. 4c). In addition, ribosome, mitochondrial translation elongation and termination, and oligosaccharyltransferase complex pathways were identified in downregulated DEGs of *Lm-*exposed male fetal brains (Fig. 4d). Analysis of the GO enrichment and KEGG analyses of upregulated DEGs in *Lm-*exposed female fetal brains demonstrated organelle related and nuclear speck pathways, whereas catenin complex and postsynaptic actin cytoskeleton pathways were identified in downregulated DEGs. (Fig. 4c and d) These findings demonstrated that placental infection had different effects on male versus female brains during neurodevelopment.

**Figure 4.**
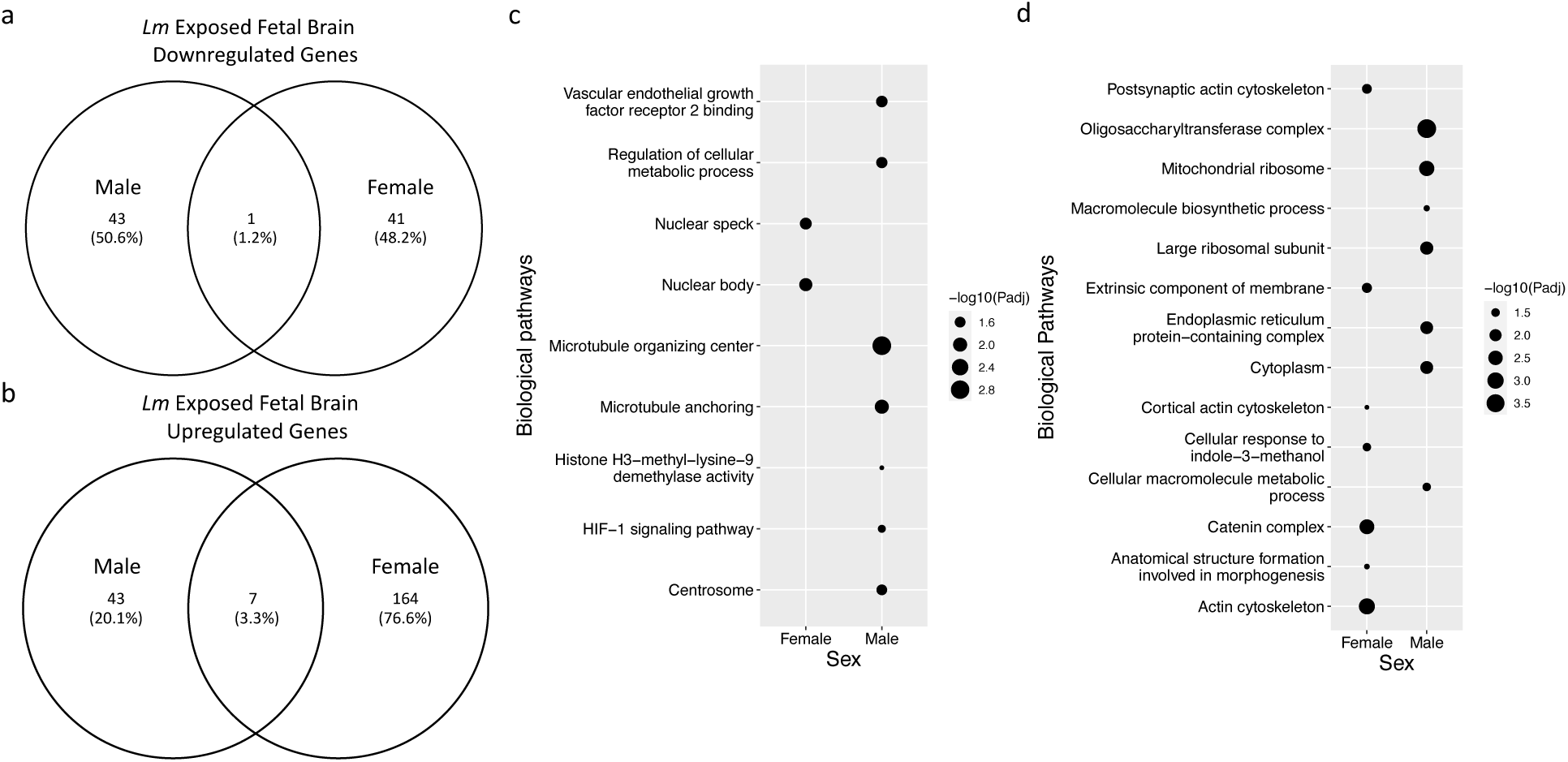
Sexually dichotomous gene expression profiles induced by placental infection. **(a, b)** Venn Diagrams representing the number of overlapping downregulated and upregulated genes in **(a)** and **(b)**, respectively. **(c, d)** Enrichment analysis of DEGs (p-adj < 0.05 of fetal brains from female and male *Lm*-exposed mice. Upregulated and downregulated biological pathways are shown in **(c)** and **(d)**, respectively.

### Placental infection induces sex-specific behavioral alterations in adult offspring

We sought to determine if altered behaviors were induced in the progeny of dams infected with *Lm*. Bacterial chorioamnionitis, which is not placental, leads to autism-like alterations in the behavioral of progeny in animals^35^, so we selected behavioral tests that are used as correlates for human ASD. For this purpose, we screened the pregnant dams with BLI to identify those with signals less than 4×10^4^ photons/sec. These mice give birth to normal-sized pups, which cannot be grossly distinguished from controls from PBS-injected dams. When the pups were 8 to 12 weeks of age, we performed behavioral assays to determine if the adult mouse offspring exhibit abnormal behavioral due to placental infection by *Lm*. We separated mice tested with behavior assays by sex to determine if placental infection results in a sex bias of these effects. First, we analyzed social interaction by using the three-chamber social approach assay to assess social impairment. Social interactions are important for forming bonds for rodents, and autism-relevant behavior mouse models have demonstrated reduction in reciprocal social interactions. *Lm-*exposed male adult offspring presented with significant reduction in social interaction time with an unfamiliar mouse, whereas *Lm-*exposed female adult offspring did not exhibit impairment in socialization (Fig. 5a; two-way ANOVA, *p* = 0.016 by Tukey’s HSD test).

**Figure 5.**
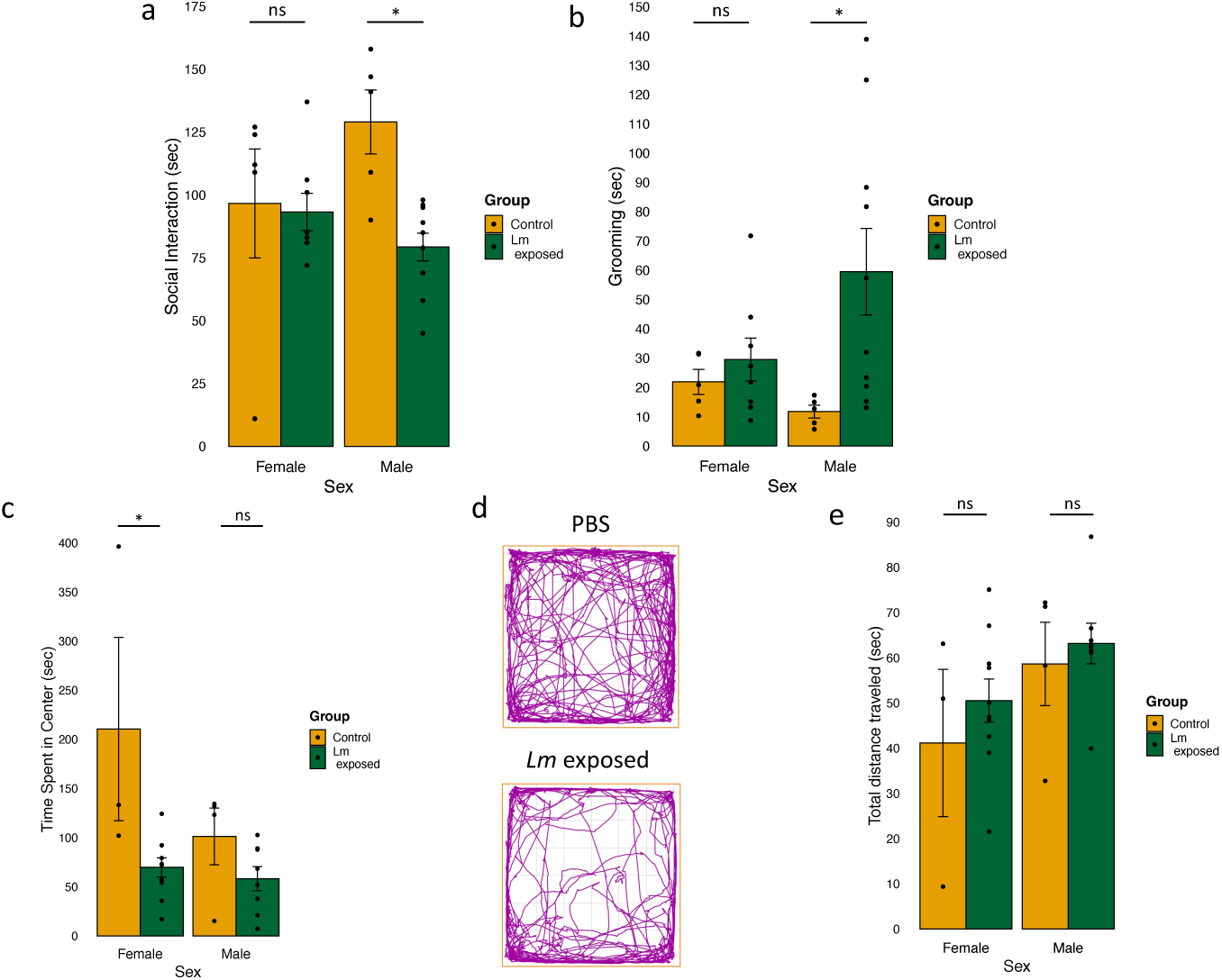
Sex-specific abnormal behaviors in the offspring of *Lm*-infected pregnant mice. **(a)** *Lm*-exposed adult male mouse offspring display deficits in social interaction (sniffing of unfamiliar mice versus inanimate objects) whereas *Lm-*exposed adult female mouse offspring show no altered behavior. **(b)** *Lm-*exposed adult male mouse offspring exhibit high levels of grooming (resembles repetitive behavior). Number (*n*) of offspring: control male (*n* = 5), *Lm-*exposed male (*n* = 10), control female (*n* = 5), and *Lm-*exposed female (*n =* 8) (a, b). **(c)** Heightened level of anxiety observed only in *Lm-*exposed adult female mouse offspring. **(d)** Differences in tracked movement during the open field exploration assay in *Lm-*exposed adult mouse offspring versus PBS controls. **(e)** No significant change in total distance traveled during open field exploration. Control male (*n* = 3), *Lm-*exposed male (*n* = 8), control female (*n* = 3), and *Lm-*exposed female (*n* = 10) used in open field exploration. Data are shown as the mean ± SEM. The behavioral assay data were analyzed by one-way analysis of variance (ANOVA) followed by Tukey’s HSD test. \**P* < 0.05.

To assess repetitive behavior with restricted interests, the duration of self-grooming behavior was examined during the three-chamber social approach assay. *Lm*-exposed male adult offspring spent significantly more time self-grooming compared to the PBS treated male mice (Fig 5b; two-way ANOVA, *p* = 0.044 by Tukey’s HSD test). However, self-grooming behavior of female adult offspring was not affected by placental infection.

Next, we examined the level of anxiety and locomotion using an open field exploration test. Rodents are hesitant to enter an unfamiliar brightly lit open field, but they gradually explore the area. Higher level of thigmotaxis, a subject remaining close to walls, is usually indicative of heightened anxiety^36^. Compared with PBS treated controls, *Lm-*exposed female adult offspring spent less time in the center of the field (two-way ANOVA, *p* = 0.011 by *post hoc* test). However, *Lm-*exposed male adult offspring did not show difference in total time spent in the center compared to the PBS treated male group (Fig. 5c). In addition, both *Lm-*exposed male and female groups showed no difference in total travel distance, which indicates locomotion activity was not affected (Fig. 5e). Together, placental infection causes abnormal behaviors in offspring that are relevant to human neuropsychiatric disorder symptoms, including elevated anxiety, increased repetitive behavior, and impaired social interaction.

### Differential expression of the activation marker c-Fos

To begin to identify changes in the adult brain due to placental infection, we immunohistochemically labeled brain sections of mice from different test groups with anti-c-Fos (Fig. 5), which has been used to characterize neuronal activity differences in MIA models^37, 38^. In these preliminary experiments, we labeled coronal sections of four male and female *Lm*-exposed mice brains and two controls mice with anti-c-Fos, a marker of brain cell activation. The results are shown in Figure 6. Sections from male mice exposed to prenatal *Lm* infection had increased c-Fos labeling compared to exposed brains from female mice and control mic. While these results are based limited in numbers of tested animals, they suggest increased neuronal activation in male mice exposed to *Lm* as a possible corollary of the sex-specific alterations of behavior

**Figure 6.**
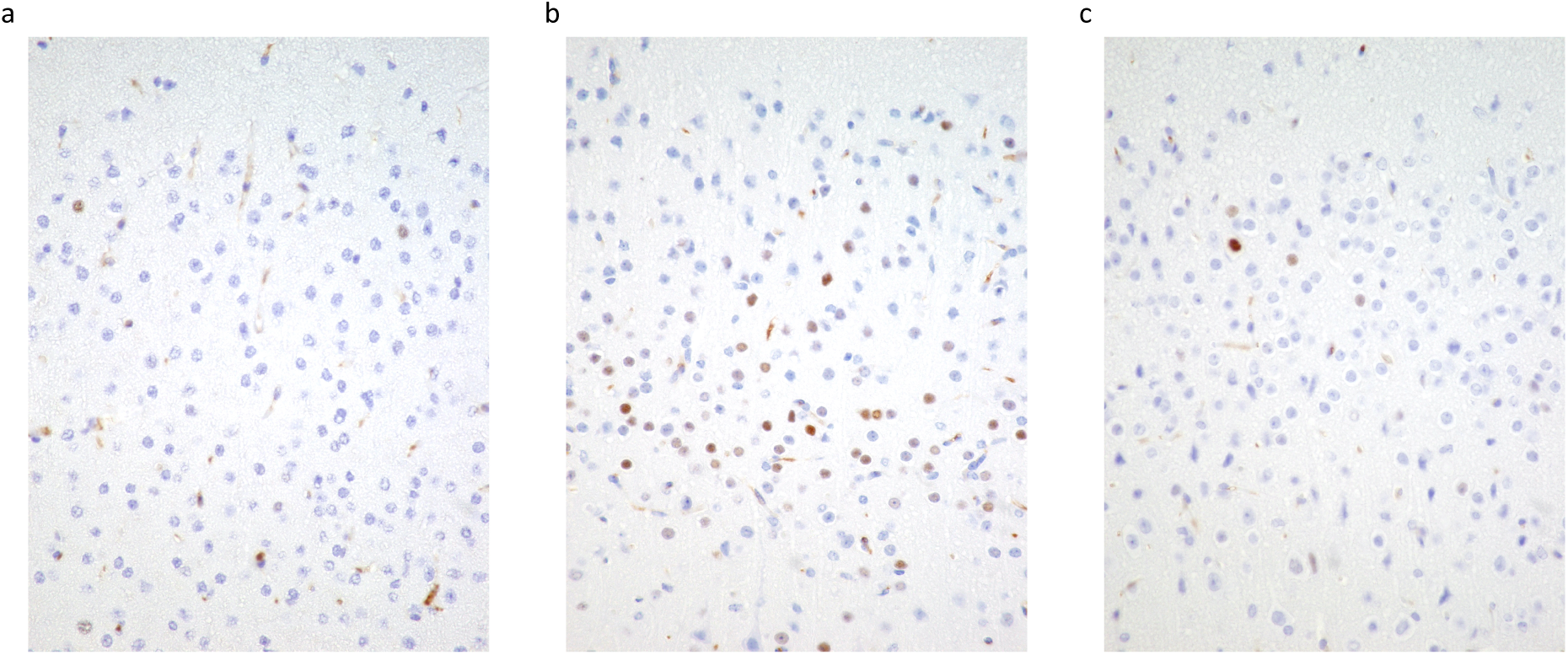
Increased nuclear c-Fos labeling in adult mouse brain due to prenatal *Lm* infection. Brains of mice that were analyzed for behavior in Figure 5 (2 infected males, 2 infected females, 2 uninfected females and one uninfected male) were immunohistochemically labeled for c-Fos. Representative sections of (a) control male; (b) *Lm*-exposed male; (c) *Lm*-exposed female. DAB chromogen (brown), hematoxylin counterstain.

## DISCUSSION

Prenatal infection is highly diverse and leads to a wide variety of outcomes both for the pregnant mother and the developing fetus. Animal models continue to reveal important mechanisms of fetal abnormality due to infection, including effects on fetal brain development that lead to abnormal behavior. Although injecting pregnant mice with immunogens, such as LPS or poly(I:C), consistently results in altered behavior and brain abnormalities in the progeny, the results of these studies are quite heterogenous^39^. In addition, these chemicals cannot be used to determine the effects of localized bacterial infections. The placental infection model using *Lm* reflects a typical subclinical infection in humans and our methods of infection allows for the analysis of abnormalities that are not due to symptomatic disease of the pregnant subject. In addition, *Lm* is well characterized and has been used for decades in placental infection models. Although listeriosis may cause serious and even fatal consequences for pregnant women and their offspring, its effect on neurodevelopment of the adult offspring has not been characterized. Here, we used bioluminescent *Lm* and the IVIS imaging system to determine if placental infection affects fetal neurodevelopment and the behavior of offspring in mice.

The identification of biological pathways in exposed whole brain transcriptome data suggests that placental infection with *Lm* dysregulates transcriptional levels of several different processes during neurodevelopment. First, the HIF-1 signaling pathway was upregulated, suggesting placental infection induces hypoxic conditions in the fetal brain during neurodevelopment. Numerous studies indicate that prenatal hypoxia results in various postnatal deficits, including reduced body and brain weight, delayed development, and impaired synaptic plasticity. Notably, *Vegfa* (vascular endothelial growth factor A), which promotes cortical interneuron proliferation, migration, and vasculature in the forebrain^40, 41^, and *Flt1* (Fms related receptor tyrosine kinase), which plays an important role in regulation of angiogenesis and development of embryonic vasculature^42, 43^, are among the main genes that are upregulated in the HIF-1 signaling pathway. Dysregulation of these genes has been identified in neuropsychiatric disorders^33, 44^. In addition, recent MIA studies demonstrated induction of hypoxia in the brain^4, 6^. Identifying elements of conservation between MIA and bacterial infection models should be examined. Interestingly, previous rodent studies have shown that prenatal hypoxia is associated with alterations in biochemical pathways during brain development, including nucleic acids process and metabolic process pathways^45, 46^. Among these pathways, *Kdm3A* (lysine demethylase 3A), which plays an important role in regulating mitochondrial biogenesis by sensing oxygen availability^47^, and *Vegfa* genes were upregulated. Lastly, protein processing in the endoplasmic reticulum (ER) pathway was downregulated due to placental infection by *Lm*. Dysregulation of protein synthesis has previously been suggested as one of the cellular responses to a hypoxic condition^48^ and implicated in various neuropsychiatric disorders^49^. Furthermore, a recent study found that poly(I:C) induced MIA triggers ER stress as a cellular response to inflammation and results in reduced protein synthesis^38^. Future work should examine the effect of placental infection on different types of cells in the fetal brain during neurodevelopment using single-cell RNA-seq.

Sexual dimorphism in neuropsychiatric disorders is well recognized. However, the basis of this dichotomy is unknown. One hypothesis proposes sex-specific vulnerability and response to environmental insults during pregnancy as one cause of sex dimorphism in these disorders. Recent MIA studies demonstrate that inflammation during pregnancy caused sex-biased placental and fetal pro-inflammatory responses^6^. Although we did not observe significant differences in BLI signals of *Lm* from placentas between male and female mice, sexually dichotomous responses are consistent with our transcriptional analysis. Interestingly, we observed more upregulated number of DEGs in brain from *Lm*-exposed female mice compared to brains of *Lm-*exposed male mice (Fig. 4b), but we did not find many biologically meaningful pathways in females. Consistent with previous MIA study, upregulation of HIF-1 signaling pathway was only enriched in *Lm*-exposed males, suggesting males are more susceptible to hypoxia during pregnancy. Future work should examine at the protein level by performing proteomic analysis to better understand how the male and female brain development is impacted by *Lm* infection during pregnancy.

Our behavioral results highlight possible pathogen specificity among rodent MIA-associated models. Injection of immune stimulants such as LPS or poly(I:C), into pregnant animals results in behavioral abnormalities in offspring that are notably relevant to ASD. Similar to these MIA-associated studies, *Lm*-exposed male offspring, but not female offspring, showed a significant reduction in social interaction and more frequent repetitive behaviors (Fig. 5a and b). These behavioral changes, and male-biased sex ratio, are observed in human ASD patients. Interestingly, we only observed significantly increased anxiety levels in *Lm*-exposed female offspring (Figure 5d), whereas MIA-associated male offspring exhibited heightened anxiety levels during open field exploration. It is important to note that numerous MIA studies have investigated behavioral changes only using male offspring^50–52^ because the prevalence of developing ASD is higher in males than in females. This difference remains to be further investigated; however, women are more likely to be diagnosed with human anxiety disorders. Another behavioral discrepancy was observed in locomotor activity. In our studies, placental infection did not alter locomotor activity in both sexes. Interestingly, Allard et al. demonstrated that prenatal infection with live Group B *Streptococcus*, a major health concern during pregnancy implicated in preterm birth and stillbirth, led to hyper-locomotor and elevated anxiety behaviors in male rat offspring, but not in female rat offspring^35^. Our contrasting results highlight the need to examine diverse prenatal pathogens, as it is becoming clear that different infections result in distinct neurological abnormalities. Studies have shown that if the locomotor activity is altered due to a treatment effect, it has a confounding effect on the movement of the animal subject during open field exploration^29^. In preliminary results, we have shown increased c-Fos labeling in brain of male but not female mice when they were exposed to *Lm* in utero (Fig. 6). This result suggests that hyperactivity of cortical neurons may be one underlying mechanism of the sexual dimorphism. Large scale and more in-depth studies will be needed to confirm this hypothesis.

One of the limitations of our studies is that individual placental BLI signals cannot be correlated with the behavior of individual offspring. Although the BLI signal of the pregnant dam can be measured using an IVIS, severity of each placental infection cannot be quantified except by sacrificing the animal. In our model, individual placentas are differentially infected by *Lm* (Fig. 1b), and our previous findings show that fetal pathologies, such as bradycardia and fetal resorption, are correlated with BLI signals from pregnant dams. Since high BLI signals in pregnant mice may result in severe postnatal consequences for the offspring, we used the animals that showed relatively low BLI signals for the behavioral analysis. We wished to compare healthy, normal-sized offspring and did not perform behavioral studies of stunted animals such as shown in Figure 1d. Another limitation of this study is the limited dose and timing of *Lm* infection we selected. According to epidemiological studies, developing psychiatric disorders is highly associated with severity and timing of the infection^53^. Furthermore, MIA-associated brain transcriptomic data from LPS and poly(I:C) have demonstrated different profiles of DEGs were observed in fetal brains that were collected at various time points^4^. Different doses and timing of *Lm* infection are likely to yield different outcomes in our behavioral and transcriptomic outcomes studies and should be performed. We have not studied the consequences of direct infection of the fetal brain, which occurs in mice with higher BLI signals, nor the effect of infection of maternal organs such as the liver or spleen, which would induce MIA. Finally, we are very interested in ascertaining the role of maternal antigen-specific immunity. These studies could be performed by vaccinating the dams before pregnancy.

In summary, we have established that placental infection by *Listeria* affects the trajectory of fetal neurodevelopment during pregnancy. We showed sex-specific dysregulation of the fetal brain transcriptome due to *Lm* infection during pregnancy. We also demonstrated that prenatal infection causes sex-specific behavioral abnormalities in offspring that resemble human ASD and anxiety-related disorders, which are known to have sexually dimorphic effects. Altogether, we have identified neurodevelopmental effects of placental infection by bacteria and expanded models of prenatal infection-associated sexual dimorphism of behavior, thus improving our understanding of the development of neuropsychiatric disorders.

## Data availability Statement

The datasets, including RNA-seq fastq and raw counts of sequencing reads, can be accessed through the NCBI Gene Expression Omnibus.

## Contributions

K.H.L. designed the experiment, performed experiments, collected data, wrote the draft and edited the manuscript. M.K and T.W. design and executed experiments and helped write the manuscript. P.P. designed and executed experiments and contributed to writing the manuscript. J. H. conceptualized the project, supervised the team with feedback and evaluation of the project, edited the manuscript. All authors read and approved the final manuscript.

## Acknowledgements

The authors would like to acknowledge Dr. Kevin Childs, the director of Genomics Core at MSU for consulting regarding the RNA-seq experiment. Additionally, we also acknowledge Dr. Alexa Veenema for the help with behavioral assays.

